# An integrative taxonomy approach reveals Saccharomyces chiloensis sp. nov. as a newly discovered species from Coastal Patagonia

**DOI:** 10.1101/2024.04.29.591617

**Authors:** Tomas A. Peña, Pablo Villarreal, Nicolas Agier, Matteo De Chiara, Tomas Barría, Kamila Urbina, Carlos A. Villarroel, Ana R. O. Santos, Carlos A. Rosa, Roberto F. Nespolo, Gianni Liti, Gilles Fischer, Francisco A. Cubillos

**Affiliations:** Facultad de Química y Biología, Departamento de Biología, Universidad de Santiago de Chile, Santiago, 9170022, Chile; Millennium Institute for Integrative Biology (iBio), Santiago, 7500574, Chile; Laboratory of Computational and Quantitative Biology, CNRS, Institut de Biologie Paris-Seine, Sorbonne Université, F-75005, Paris, France; Université Côte d’Azur, CNRS, INSERM, IRCAN, Nice, France; Millenium Nucleus of Patagonian Limit of Life (LiLi), Santiago, 7500574, Chile; Centro de Biotecnología de los Recursos Naturales (CENBio), Facultad de Ciencias Agrarias y Forestales, Universidad Católica del Maule, Talca, Chile; Departamento de Microbiologia, ICB, C.P. 486, Universidade Federal de Minas Gerais, Belo Horizonte, MG, 31270-901, Brazil; Instituto de Ciencias Ambientales y Evolutivas, Universidad Austral de Chile, Valdivia, Chile; Center of Applied Ecology and Sustainability (CAPES), Facultad de Ciencias Biológicas, Universidad Católica de Chile, Santiago, Chile

**Keywords:** *Saccharomyces chiloensis* sp. nov, Ortho Average Nucleotide Identity (OANI), Patagonia, integrative taxonomy, yeast

## Abstract

Species delineation in microorganisms is challenging due to the limited markers available for accurate species assignment. Here, we applied an integrative taxonomy approach, combining extensive sampling, whole-genome sequence-based classification, phenotypic profiling, and assessment of interspecific reproductive isolation. Our work reveals the presence of a distinct *Saccharomyces* lineage in *Nothofagus* forests of coastal Patagonia. This lineage, designated *Saccharomyces chiloensis* sp. nov., exhibits 7% genetic divergence from its sister species *S. uvarum*, as revealed by whole-genome sequencing and population analyses. The South America-C (SA-C) coastal Patagonia population forms a unique clade closely related to a previously described divergent *S. uvarum* population from Oceania (AUS, found in Australia and New Zealand). Our species reclassification is supported by a low Ortho Average Nucleotide Identity (OANI) of 93% in SA-C and AUS relative to *S. uvarum*, which falls below the suggested species delineation threshold of 95%, indicating an independent evolutionary lineage. Hybrid spore viability assessment provided compelling evidence that SA-C and AUS are reproductively isolated from *S. uvarum*. In addition, we found unique structural variants between *S. chiloensis* sp. nov. lineages, including large-scale chromosomal translocations and inversions, together with a distinct phenotypic profile, emphasizing their intraspecies genetic distinctiveness. We suggest that *S. chiloensis* sp. nov diverged from *S. uvarum* in allopatry due to glaciation, followed by post-glacial dispersal, resulting in distinct lineages on opposite sides of the Pacific Ocean. The discovery of *S. chiloensis* sp. nov. illustrates the uniqueness of Patagonia’s coastal biodiversity and underscores the importance of adopting an integrative taxonomic approach in species delineation to unveil cryptic microbial species. The holotype of *S. chiloensis* sp. nov. is CBS 18620^T^.

## INTRODUCTION

The concept of a species is a highly debated term in biology, especially when applied to microorganisms^1^. Species delineation, the process of defining and distinguishing species within a taxonomic group, faces important challenges in fungi and yeast, mostly because the species’ morphological concept does not apply^2^. Over time, advances in molecular biology and genomic analysis have significantly driven the identification of different species in eukaryotic microorganisms, regularly based on the phylogenetic and biological species concept^1,3,4^. In this sense, yeasts with a known sexual cycle can be assessed regarding reproductive isolation, following the biological species concept^5^. At the genetic level, most yeast species can be identified based on sequence divergence in the more rapidly evolving internal transcribed spacer (ITS), as well as the slowly evolving SSU (Small Subunit) and LSU (Large Subunit) rRNA genes. Hence, it is accepted that nucleotide divergence exceeding 1% in the D1/D2 domain of the LSU rRNA gene suggests distinct species^6,7^. However, the use of these markers may be influenced by population genetics and demographic processes, especially in temperate and arctic regions. In these regions, glaciation processes in the past have driven some species and sub-populations into allopatric distributions during colder periods and sympatric distributions during warmer periods^8–10^. Phases of allopatry and post-glacial range expansions can manifest as intricate genetic structures within and between species. For instance, while closely related species may share identical ITS and D1/D2 sequences they can show considerable divergence elsewhere in the genome^11,12^. Thus, using a few genetic markers alone can lead to overly conservative species delineation^7^.

Recent advances have enabled the use of whole-genome data for species delineation, which is essential for comparing pairwise average nucleotide identity (ANI) between species across the entire genome. ANI quantifies the DNA-level similarity between two strains, correlating with DNA-DNA hybridization. ANI values exceeding 95 ± 0.5% typically suggest conspecificity^13–15^. Recently, Lachance et al. (2020) suggested in a large data set on haplontic, heterothallic *Metschnikowia* species that ANI of 95% constitutes a good threshold for species recognition, while values lower than 95% (ANI’D>D5%) provided evidence that speciation has taken place^16^. Recently, long-read sequencing technologies have significantly improved genome annotations and ANI estimations by only considering symmetrical nucleotide identity comparisons between ortholog pairs, known as average nucleotide identity by orthology (OrthoANI) ^17^. Therefore, a widely accepted consensus delineation threshold for yeast species is an OANI value of 95%^16^. However, delineating yeast species still requires careful consideration of all available data, including genetic variation, phenotypic divergence, ecological niches, and population history. A general current view is to accommodate the so-called ‘integrative taxonomy’ approach, incorporating genomic, reproductive, and morphological characters for species delineation^18,19^. The concept of integrative taxonomy acknowledges the influence of ecological and evolutionary processes on species boundaries, providing a robust approach to defining new species ^19^.

The *Saccharomyces* genus is the most prominent and well-studied group across the Saccharomycotina subphylum. This genus includes eight species, many of them with significant economic and scientific importance^5^. Recent advances benefiting from bioprospecting efforts and using genomics and phylogenetics tools have revealed two additions to the *Saccharomyces* clade: *Saccharomyces eubayanus* and *Saccharomyces jurei*^20–22^. *S. eubayanus*, first identified in the temperate forests of Patagonia, coexists sympatrically with *S. uvarum*^23–25^. Despite their taxonomic closeness, in-depth genomic analyses have revealed substantial genetic diversity and evolutionary divergence between these yeast species^23,26^. Comparative genomic studies have emphasized differences in chromosomal structure and gene content, highlighting their genetic distinctiveness^26,27^. The genetic relationship between *S. eubayanus* and *S. uvarum* is characterized by evidence of shared ancestry and common glaciation periods in Patagonia, hinting at a shared evolutionary origin within the Patagonian region approximately 16 MYA^28,29^. During these periods, vast ice sheets covered a substantial portion of the land in southern Patagonia, including some regions of the southern Pacific islands. This was succeeded by an extensive floristic recolonization and ecological succession during warmer periods^30,31^. These events probably played a vital role in shaping the current genetic structure of *Saccharomyces* species in Patagonia ^23^.

*S. uvarum* populations in the Patagonian Andes contain significant genetic variation, more than anywhere else in the world, and they have diverged into two major sub-groups: South America SA-A and SA-B^28,32^. Intriguingly, Holarctic (HOL) strains from Europe and North America clustered alongside the SA-A strain, indicating possible migration patterns in both directions between Europe and America^32^. Intriguingly, a vastly divergent lineage of *S. uvarum* (AUS) has been identified in Tasmania ^28^ and New Zealand^33^, forming a distinct branch within this yeast species. The AUS lineage represents an early diverging cluster and demonstrates partial reproductive isolation from other lineages^28^. Although most *S. uvarum* populations are found in Patagonia, there have been no reports of the presence of the AUS lineage in South America, nor reports of *Saccharomyces* isolation efforts in Coastal Patagonia or nearby islands with similar geography. The factors contributing to the significant genetic divergence in AUS could be linked to an allopatric speciation process due to geographical separation, underscoring the complexity of the species and the potential for isolated populations to evolve distinct genetic and phenotypic patterns.

In this study, we assessed the diversity of *Saccharomyces* lineages in Coastal Patagonia and discovered a novel *Saccharomyces* lineage isolated from *Nothofagus dombeyi* forests on islands in Southern Chile. Phylogenomic analysis revealed a close relationship with the AUS population previously described in *S. uvarum*. We applied an integrative taxonomy approach that included phylogenomic assessments, postzygotic reproductive isolation evaluations, and phenotypic characterization. The new lineage displayed considerable nucleotide divergence, reproductive isolation, and a range of structural variants, including notable large translocations found to be polymorphic between Chilean and Australian populations. Our data suggests that the Chilean and Australian populations originated in allopatry. As such, we propose recognizing them as a distinct species named *Saccharomyces chiloensis* sp. nov.

## METHODS

### Yeast isolation

Samples were collected and processed as previously reported^23,34^ from bark samples from 106 Coigüe *(Nothofagus dombeyi)* trees from three different localities in coastal Patagonia: Chiloé, Reserva Costera Valdiviana, and San Jose de la Mariquina (**Table S1a**). *Saccharomyces* identification was performed as previously reported^23^. First, the internal transcribed spacer (ITS) was amplified and sequenced using primers ITS1 and ITS4^35^ to discriminate between *Saccharomyces* and non-*Saccharomyces* colonies. Subsequently, differentiation between *Saccharomyces* species was performed using the polymorphic marker *RIP1* by amplification, enzyme restriction, and Sanger sequencing as previously reported^23,25^. Furthermore, we used 42 strains reported as *S. uvarum* from our yeast collection^23^ to reveal the complete diversity landscape of *Saccharomyces* isolates from Patagonia (**Table S1b**). Representative sequences for *ITS* and *RIP1* can be found under the code PP618688-PP618690 and PP578963-PP578965.

### Whole-genome sequencing and Variant Calling

Genomic DNA was extracted using the Qiagen Genomic-tip 20/G kit (Qiagen, Hilden, Germany) as previously described ^23^ and sequenced using an Illumina NextSeq 500 system. Raw reads quality was checked using FastQC^36^. Reads were processed with fastp 0.23.2 (-q 20 -F 15 -f 15)^37^. Trimmed reads were aligned against the *S. uvarum* CBS7001^T^ ^38^ using BWA-mem (option: -M -R) ^39^. Mapping quality and summary statistics were obtained using Qualimap v.2.2.2-dev ^40^. SAM files were sorted and transformed to BAM format using Samtools 1.14^41^. BAM files were tagged for duplicates using MarkDuplicates of Picard tools 2.27.2^42^. Variant calling and filtering were done using GATK 4.2.3.0^43^. Specifically, variants were called per chromosome per sample using HaplotypeCaller (default settings). Next, a variant database was built using GenomicsDBImport. Genotypes per chromosome were called using GenotypeGVCFs (-G StandardAnnotation). Genome-wide genotype for each sample was merged using MergeVcfs. We applied recommended filters for coverage (>10 mapping reads = “FORMAT/DP>10”) and quality (–minQ 30). Multisample VCF was further filtered when needed with vcftools 0.1.16. We only considered SNPs without missing data using –max-missing 1^43^. Statistics from the final VCF file were obtained using bcftools stats command^41^.

To obtain individual consensus genomes for each strain, we used the VCF data together with the CBS7001^T^ genome. First, we used the vcf-subset mode of vcftools 0.1.16 to extract individual SNPs from each strain. Then, we indexed each VCF with IndexFeature mode from GATK 4.2.3.0. Finally, using FastaAlternateReferenceMaker from GATK 4.2.3.0 we obtained consensus fasta genome per strain.

### Phylogenomic reconstruction

To perform the phylogenomic analysis using our biallelic SNP dataset, we transformed a VCF file containing 1,504,923 SNPs into phylip format using vcf2phylip^44^. This was used as input for the IQ-TREE^45^ to generate a maximum likelihood phylogeny with the ultrafast bootstrap option and ascertain bias correction (-st DNA -o CL815 -m GTR+ASC -nt 8 -bb 1000)^46^. The number of parsimony informative sites was 833,534. Trees were visualized on the iTOL website^47^.

Phylogenomic analysis using orthologues genes was done using OrthoFinder V2.5.5^48^. Specifically, the complete set of ORFs (in peptide sequence) was extracted from the Augustus output^49^, and then used as input for OrthoFinder V2.5.5^48^. Visualization of the tree was done with iTOL website^47^.

### Population genetic analyses

First, a VCF file’s thinned version was generated with vcftools 0.1.16 (--thin 500)^41^. Ancestry estimation was explored by running ADMIXTURE^50^ using k=2 to k=10 five times per k using different seed numbers. Cross-validation errors of each run were used to choose the best number of populations. Results were visualized using Pong^51^. Additionally, to perform a clustering analysis using SMARTPCA^52^, we kept only those individuals with no evidence of admixture in the ADMIXTURE analysis. Using the same data set of phased individuals with Beagle 3.0.4^53^, we performed a fineSTRUCTURE analysis as previously reported^54^ considering a constant recombination rate between consecutive SNPs based on the *S. cerevisiae* average recombination rate (0.4 cM/kbp)^55^. Pairwise nucleotide diversity (*π*) and Fixiation index (*F*_st_) were estimated using the R package PopGenome^56^. To predict the number of generations since the most recent common ancestor of any pair of lineages we used the mutation parameter^57^, ******, in which ****** represents the number of generations. Considering that the single-base mutation has been reported similar for multiples yeast species^58,59^, we used the mutation rate µ = 1.84 x 10^-10^ (mutations per generation) previously reported in *S. cerevisiae* for calculations^60^.

### Species delineation analysis

To delineate species boundaries based on genetic differences, we used the average nucleotide identity (ANI) as a main criterion^61^. All ANI values were calculated against the *S. uvarum* reference genome, CBS7001^T^, by using OrthoANI algorithm^17^. As described for yeast, we used the 95-96% ANI value as cut off for species delinitation^15,16^.

### Spore viability analysis of interpopulation hybrids

To test spore viability between crosses from different lineages, diploid cells were sporulated on 2% potassium acetate agar plates (2% agar) for at least 7 days at 25 °C. Meiotic segregants were obtained by dissecting tetrad ascospores treated with 10 µL Zymolyase 100T (50 mg/mL) on YPD agar plates with a SporePlay micromanipulator. Spores were crossed against the *S. uvarum* haploid CBS7001^T^ strain derivate (HOL, *HO::NatMx*)^62^. Hybrid candidates were picked from selective media^62^ and confirmed by PCR amplification of the *MAT* locus (hybrids were *Mat-a*/*Mat-α, HO::NatMx*). The hybrids were then sporulated as previously mentioned. At least 96 ascospores per hybrid were dissected in YPD plates and incubated at 25 °C for two days. Spore viability was calculated using the formula (n° of viable spores/n° of dissected spores)D×D100^21,63^. Additionally, spores from hybrids with low reproductive success were sporulated and dissected for a F2 collection.

### Nanopore sequencing, de novo assembly and genome annotation

For nanopore sequencing EXP-NBD104 barcodes and SQK-LSK109 adapters were ligated, and library was loaded on R9.4.1 flowcells; sequencing was performed on a MiniIon device (Nanopore, Oxford, UK). Raw fast5files were converted for fastq files using Guppy, followed by Porechop for the removal of adapters and barcodes^64^. Canu v2.2^65^ with default setting was used for assembly, followed by two rounds of both racon v1.4.3^66^ and pilon v1.2.4^67^ for long and short read polishing. Genome assemblies were annotated using the pipeline LRSDAY^68^ using the complete genome of the CBS7001T type strain as training input for AUGUSTUS^49^. Gene function was predicted using as reference the *S. cerevisiae* S288c genome, all the other parameters were set as default. Genome completeness was assessed by using BUSCO 5.5.0^69^. Assemblies were compared with CBS7001^T^ using nucmer^70^, and structural variants (SVs) were called using MUM&Co^71^. Calls on the ribosomal DNA were not considered. Divergence was re-estimated from whole genome alignment by using Mummer4 dnadiff command^70^.

### Introgression identification

Introgressions from *S. uvarum* were identified using a combination of SNP density analysis and mapping mean coverage within 1 kb non-overlapping windows^28^. To achieve this, we mapped one representative strain from each *S. uvarum* lineage to a *de novo* assembly and called variants as previously described^23^. Regions exhibiting zero SNP density and more than 20x mapping coverage were selected as potential introgressions.

### Phenotypic diversity among isolates

The different isolates were subjected to morphological, physiological, and biochemical characterization under solid media using the replica-plating technique. This physiological and biochemical characterization was done using the standard methods described by Kurtzman et al., with a few modifications^72^. The growth of the yeast strains was tested on YNB plates containing 2% of each carbon source. For cell morphology analysis, observations were made using a Zeiss AXIO Imager M2 microscope equipped with differential interference contrast optics (Axiocam 506 mono) after 3 days of growth in YPD broth, incubated at 25°C. Ascospore production was conducted on 2% potassium acetate agar plates (2% agar) for at least 7 days at 25 °C. Pseudohyphae and true hyphae formation were detected using the Dalmau plate culture method, as described by Kurtzman et al^72^.

To determine the phenotypic diversity of the different isolates, we estimate the growth and biomass production in several environmental conditions that are representative of different yeast habitats, either in nature or in industrial settings, as previously described^73^. For this, we performed a high-throughput phenotyping assay in 96-well microculture plates. Briefly, cells were pre-cultivated in 200 μL of a YNB medium supplemented with 2% glucose for 48h at 25°C. Then, strains were inoculated to an optical density (OD) of 0.03–0.1 (wavelength 620 nm) in 200 μL of media and incubated without agitation at 20°C for 24 h (YNB control) and 48-96 h under other conditions. OD was measured every 30 min using a 620 nm filter. Each experiment was performed in triplicates. The conditions considered were Yeast Nitrogen Base (YNB) supplemented with 2% at 25 °C; 2% glycerol, 2% glucose 2% NaCl; 2% maltose; 2% Fructose; 2% Sucrose; 2% Galactose and 2% glucose + 5% ethanol. Area under the curve (AUC) was estimated with the R package growthcurver^74^.

### Statistical analysis

All statistical analyses were performed using biological triplicates. The difference was considered statistically significant at *p*-value < 0,05. The comparison of The Area Under the Curve (AUC) between strains/lineages was performed using a one-way ANOVA. The heat map was created using R package, ComplexHeatmap^75^.

### Data availability

Fastq sequences and the type strain genome annotation were deposited in the National Center for Biotechnology Information (NCBI) as a Sequence Read Archive under the BioProject accession number PRJNA1084000. The scripts and codes used in our study were deposited in Github: https://github.com/TomaasP/SchiloGenomics/tree/main.

## RESULTS

### A highly divergent *Saccharomyces* lineage discovered in coastal Patagonia

To identify the *Saccharomyces* lineages in the Coastal Patagonian region^23,73^, we conducted a survey targeting *N. dombeyi* forests along the South Chilean Pacific coast (**Figure 1A**). We collected bark samples from different sampling points in the Chiloe island and the Valdivian coastal area, isolating 106 *Saccharomyces* strains (**Table S1a**). Subsequently, we performed genotyping on these isolates by sequencing the highly polymorphic *RIP1* marker, enabling the discrimination between *S. uvarum* and *S. eubayanus* species. Interestingly, our genotyping assay unveiled three distinct genotypes for the *RIP1* marker: two patterns corresponding to *S. uvarum* and *S. eubayanus*, while a third, matched to a high divergent *S. uvarum* lineage (AUS from Almeida 2014), identified in 43 isolates (**Figure S1A, Table S1a**). Further Sanger sequencing of these isolates’ ITS, LSU, and SSU regions confirmed a 100% sequence identity with *S. uvarum*, strongly suggesting that these isolates likely belong to this species (**Figure S1B and C, Table S1a**).

**Figure 1.**
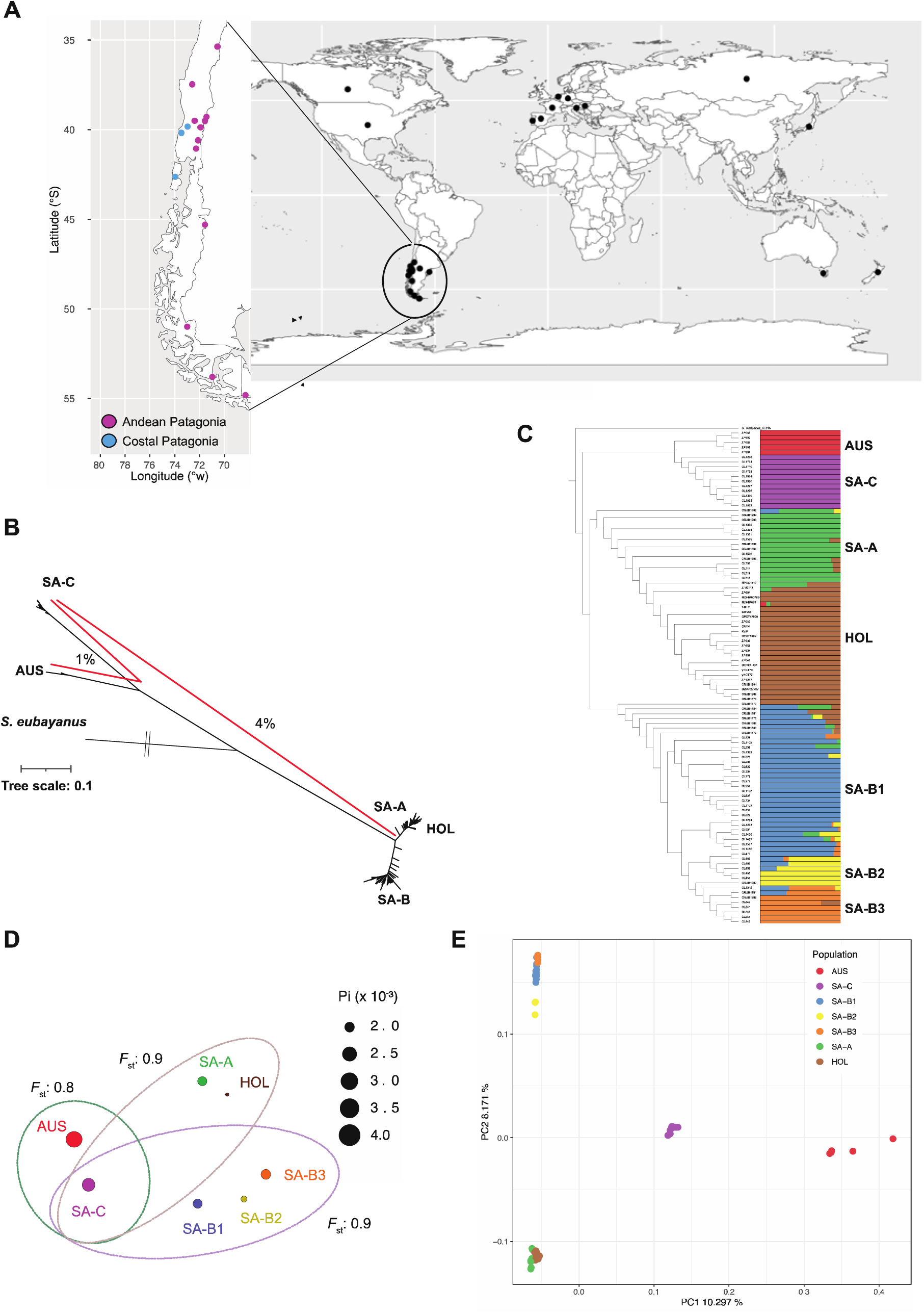
Geographic distribution and population structure of Coastal Patagonian isolates. (A) World map illustrating the distribution of sequenced genomes of *S. uvarum* globally (black circles), with specific locations in Chile in Andean (purple) and Coastal (blue) Patagonia where new strains were isolated. (B) Unrooted maximum likelihood tree inferred from an alignment of 100 strains. Red lines indicate the average nucleotide distance between lineages using Illumina reads. The tree contains five main lineages: South America (SA-A, SA-B and SA-C), Australia (AUS) and Holarctic (HOL). Tree scale is substitutions per site (total SNPs across the whole genome size). (C) Non-scaled Maximum likelihood (ML) phylogenetic tree and population analysis using ADMIXTURE. The optimum of k=7 groups is shown. Lineages identified are: South America (SA-A (green), SA-B1 (blue), SA-B2 (yellow), SA-B3 (orange) and SA-C (purple)), Australia (AUS, red) and Holarctic (HOL, brown). *S. eubayanus* CL815 was used as outgroup. (D) Population differentiation (*F*_ST_) and nucleotide diversity (π) of the seven lineages. Circle size indicates π value. Dashed circles represent the *F*_ST_ value between groups. (E) Distribution of genomic variation in 100 strains based on the first two components from a PCA analysis. Color codes match the clusters presented in Panel C. Each dot on the graph represents an individual strain.

To unravel the phylogenetic relationships of the coastal Patagonia strains within the *S. uvarum* phylogeny, we obtained short-read whole-genome sequences from 19 isolates representing various localities in Coastal Patagonia. Additionally, we sequenced 37 *S. uvarum* isolates previously obtained by our group from diverse locations in Andean Patagonia (**Table S1b, Figure 1A**), and we included 43 *S. uvarum* genomes from previous reports (**Table S1c**)^28,76^, along with one representative *S. eubayanus* isolate (**Table S1b**)^23^. In total, we analyzed 100 strains: 17 strains isolated from Argentina, 56 from Chile, 5 from Australia, and 19 from the Holarctic regions (Europe, North America, and Asia) (**Table S2a and S2b**). The maximum-likelihood phylogenetic tree, employing *S. eubayanus* as an outgroup, reaffirmed earlier observations: the AUS isolates exhibited the greatest genetic divergence from the SA-A and SA-B South American populations (**Figure 1B**). Notably, the isolates from coastal Patagonia formed a distinct lineage, closely clustering with AUS, strongly suggesting the presence of a unique independent lineage.

We further utilized the ADMIXTURE software to delve into the population structure of *Saccharomyces* isolates, using from k 2 to 10 (**Figure S2**). The analysis suggests 7 populations as the best-fitting model (**Figure S3**), which revealed distinct lineages predominantly corresponding to the geographical origin of the isolates. This analysis sheds light on a different lineage encompassing the coastal Patagonian isolates (hereafter termed SA-C), closely clustering with the AUS population. Moreover, multiple *S. uvarum* lineages were identified within South America (SA-B1, SA-B2, and SA-B3) (**Figure 1C**). This topology was consistently supported by both *F*_st_ and PCA analyses (**Figure 1D and E, Table S3),** demonstrating substantial genetic differentiation between the AUS and SA-C lineages compared to the SA-A and SA-B Andean Patagonian lineages. FineSTRUCTURE analysis echoed these findings, highlighting an equivalent number of distinct lineages (**Figure S4**). Estimating genetic diversity (π) revealed in the SA-C and AUS populations a mean π = 0.0032 and 0.0042, respectively, surpassing the genetic diversity observed in other *S. uvarum* populations (**Figure 1D, Table S4**). Patterns of pairwise nucleotide variation using Illumina reads showed that AUS and SA-C were, on average 3.69% and 4.06 % divergent from the *S. uvarum* reference genome (CBS 7001^T^, HOL), respectively (**Table S5, Figure S5**). Interestingly, the average pairwise nucleotide divergence between the AUS and SA-C lineages was, on average, 1.03%, suggesting a genetic differentiation between these two lineages (**Table S5**). The high genetic divergence between AUS and SA-C relative to *S. uvarum* suggests that these groups of strains could be reproductively isolated and represent a distinct evolutionary lineage.

### Low spore viability between SA-C and other *S. uvarum* lineages

To assess the impact of nucleotide sequence divergence on offspring viability, we crossed spores from SA-C and AUS isolates with the *S. uvarum* CBS7001^T^ reference strain, estimating spore viability. Crosses involving representative SA-C and AUS strains (CL1604 and ZP966 strains, respectively) exhibited markedly lower spore viability (6.02% and 5.09%, respectively, **Table S6, Figure 2**). The resulting spores were sterile, and several did not undergo tetrad formation. On the contrary, crosses between representative strains from different *S. uvarum* populations with CBS7001^T^ had higher spore viability, ranging between 50% and 81%, consistent with their lower genetic divergence (**Table S6, Figure 2**). This reproductive isolation suggests distinct barriers within the SA-C and AUS lineages compared to other *S. uvarum* populations.

**Figure 2.**
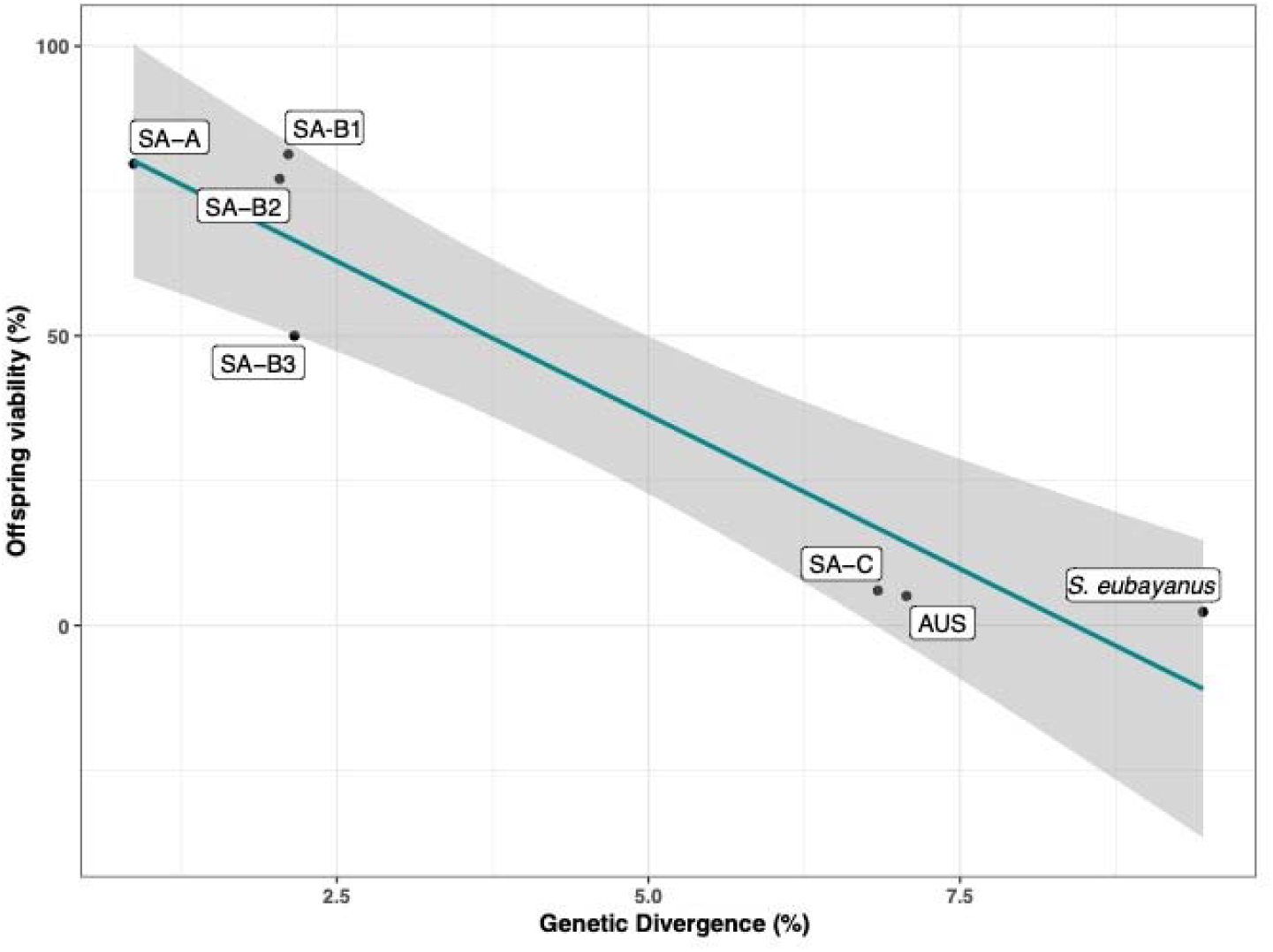
Offspring viability against *S. uvarum*. Plot depicting spore viability between lineages and genetic divergence (measured as nucleotide identity). Each dot compares one representative strain mated against the *S. uvarum* reference strain CBS7001^T^. A linear regression model was applied, with an Adjusted R-squared of 0.8891 and a *p*-value of 0.001. 95% confidence intervals are in gray.

We crossed representative SA-C and AUS strains (CL1604 and ZP966 strains, respectively) to explore potential lineage relationships between SA-C and AUS, revealing a 25.7% spore viability (**Table S6**). This indicates genetic differences beyond nucleotide divergence, likely involving additional genetic variants, such as structural variants (SVs). Interestingly, diploid SA-C and AUS strains exhibited normal spore viabilities, demonstrating that they are homothallic and ruling out inherent reproductive issues (**Table S6**). These results suggest reproductive isolation between SA-C, AUS, and other *S. uvarum* populations, possibly due to nucleotide divergence. Additionally, SA-C and AUS lineages may exhibit additional genetic variants impacting their reproductive status. Considering the nucleotide divergence and reproductive isolation, we propose classifying the SA-C and AUS lineages as a distinct species, named ***Saccharomyces chiloensis* sp. nov.** for their most frequent isolation in the Chiloe island.

### Coastal Patagonian lineages exhibit large genomic rearrangements

To identify SVs that could shed light on the low spore viability between the SA-C and AUS *S. chiloensis* sp. nov lineages, we obtained *de novo* assemblies using ONT long-read sequences from a subset of 5 representative strains from both *S. chiloensis* sp. nov lineages (**Table S7**). Using the assemblies, we identified SVs with Mum&Co of at least 50 bp relative to the *S. uvarum* and *S. eubayanus* reference genomes (CBS7001^T^ and CBS12357^T^, respectively). To ensure accuracy and avoid false positives stemming from assembly errors, we filtered the dataset for SVs size (> 1kb), resulting in the identification, on average, of 72,6 and 85,8 SVs at the lineage level between the SA-C *S. chiloensis* sp. nov. lineage and *S. eubayanus* and *S. uvarum*, respectively (**Figure S6, Table S8**). Interestingly, we identified five large rearrangements across all SA-C *S. chiloensis* sp. nov. strains: one reciprocal translocation (ChrVI-ChrX^t^, ChrX-ChrVI^t^), two non-reciprocal translocation between three chromosomes (ChrXV-ChrII^t^ and ChrII-ChrVII^t^), one inversion (ChrXIII^I^) and a deletion on the right arm of ChrVII relative to *S. uvarum* (**Figure 3A**). The translocated segments spanned approximately 54 (ChrVI-ChrX^t^), 318 (ChrX-ChrVI^t^), 433 (ChrXV-ChrII_t_) and 143 (ChrII-ChrVII^t^) kb, and comprised 19, 145, 212, and 64 genes, respectively. The inversion spanned approximately 487 kb, covering 225 genes (**Table S9**). A similar comparison between the AUS lineage and *S. uvarum* revealed the presence of the reciprocal translocation previously described between ChrVI-ChrX^t^ (**Figure 3B**), indicating that this reciprocal translocation would be shared across *S. chiloensis* sp. nov. However, not all SVs are conserved across *S. chiloensis* sp. nov lineages (**Figure 3C**). In this sense, the two non-reciprocal translocations (ChrXV-ChrII^t^ and ChrII-ChrVII^t^) along with the ChrXIII inversion and Chrm VII deletion are exclusively found in the SA-C lineage, likely arisen following the split of the AUS and SA-C lineages along the SA-C branch (**Figure 3C**, **Table S9**).

**Figure 3.**
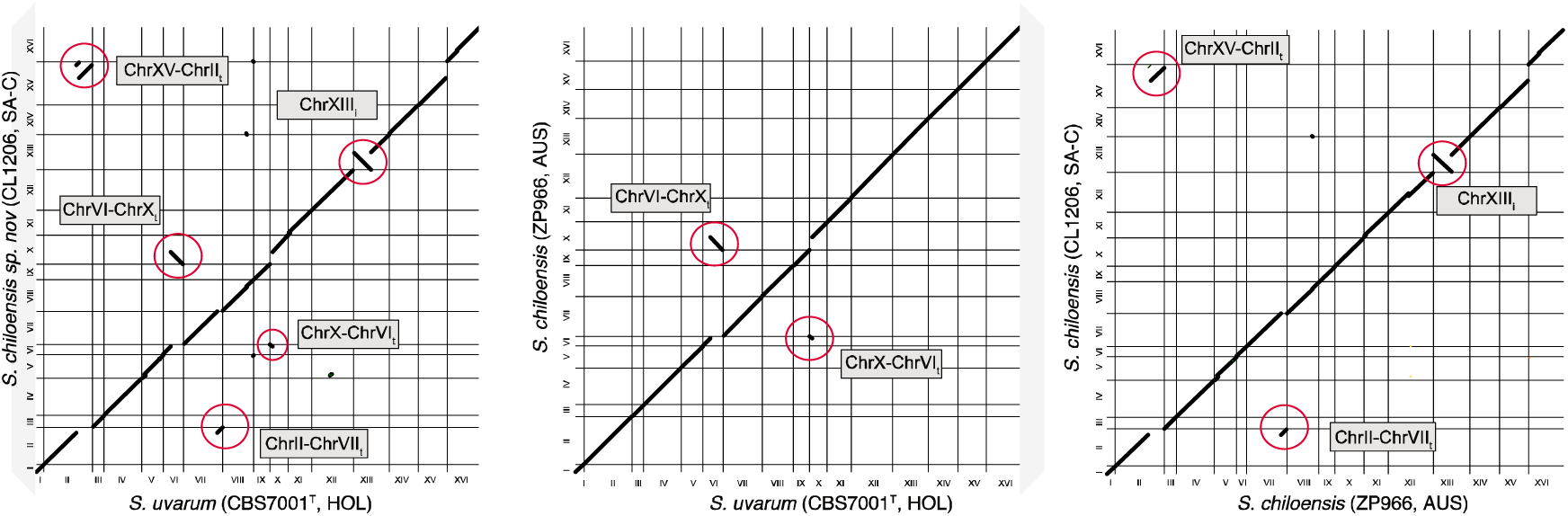
Structural variants identification between *Saccharomyces chiloensis* sp. nov. and other *Saccharomyces* species. (A) Genome synteny analysis of the S*. uvarum* reference strain CBS7001^T^, and (A) *S. chiloensis* sp. nov. CL1206 strain (SA-C) and (B) ZP966 strain (AUS). The dot plot representation depicts the DNA sequence identity between the genomes. Translocation and inversions are highlighted in red circles and grey rectangles indicate the chromosomes involved. Translocations were mapped in all the *S. chiloensis* sp. nov. strains sequenced in this study. (C) Genome synteny analysis of the *S. chiloensis* sp. nov. CL1206 strain (SA-C) and ZP966 strain (AUS) strains.

Since many genes are affected by SVs, we decided to explore how gene content varies between *Saccharomyces* species. We first annotated all the *de novo* assembled genomes and then constructed a phylogeny considering both *S. chiloensis* sp. nov lineages and the different *Saccharomyces* reference genomes (**Figure 4A**). Interestingly, we reveal the apparent greater genetic distance between SA-C and AUS relative to *S. uvarum* compared to other lineages within the same species (**Figure 4A, Table S10**). We used the Ortho Average Nucleotide Identity (OANI) tool to explore the mean nucleotide identity between *S. chiloensis* sp. nov and *S. uvarum* lineages. The SA-C vs. *S. uvarum* comparison showed 92.9% identity, while the AUS vs. *S. uvarum* comparison showed 93.3%. These are lower values than any of the other comparisons between the remaining *S. uvarum* lineages revealed (**Figure 4B**, **Table S10**). When we performed a similar analysis using highly divergent lineages in *S. paradoxus*^77^ and *S. kudriavzevii* (nucleotide divergence ∼ 4.7%), the comparisons denoted values near the 95% species delineation limit (**Table S10**), demonstrating that the SA-C and AUS lineages are more divergent than others in the genus (**Figure 4C**).

**Figure 4.**
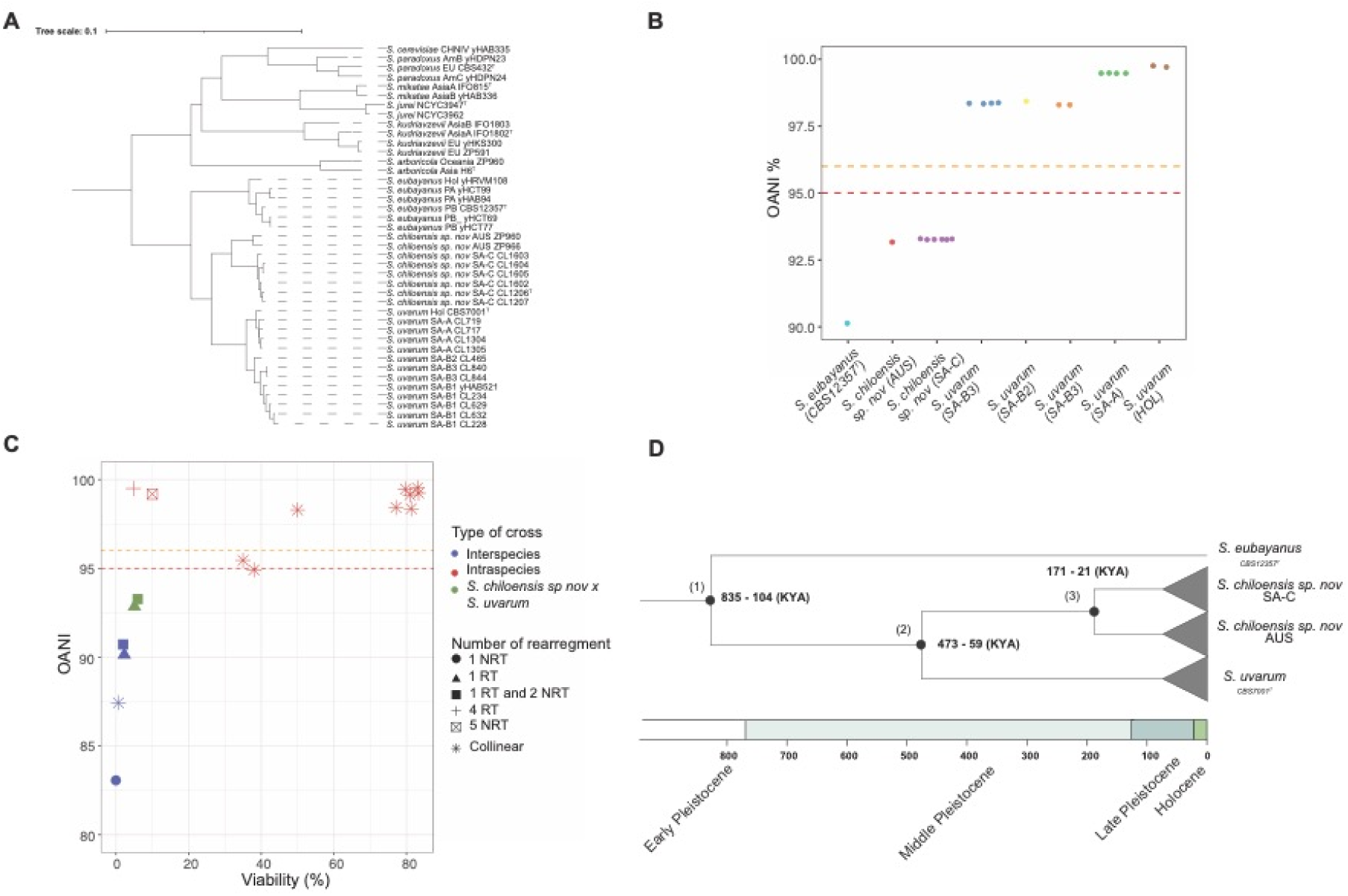
Phylogenetic relationship across different *Saccharomyces* species. (A) Maximum likelihood (ML) phylogenetic tree using different *Saccharomyces* species and lineages where long-reads are available. The tree was built considering 5,772 orthologues genes using OrthoFinder. Branch lengths represent the average number of substitutions per site (tree scale). Representative lineages from each species were selected, and annotated genomes were obtained from previous studies using publicly available data. (B) OANI values were estimated using each genome from the ONT assemblies across the different lineages against *S. uvarum* reference genome (CBST7001^T^). Dashed red and orange lines denote the 95% and 96% species delineation thresholds. (C) Plot depicting OANI values and offspring viabilities comparing different sister *Saccharomyces* species (blue dots: *S. cerevisiae* x *S. paradoxus*^55,78^; *S. jurei* x *S. mikatae*^21^ and *S. eubayanus* x *S. uvarum*), *Saccharomyces* lineages within the same species (red dots, intraspecies crosses in *S. cerevisiae*^78^, *S. kudriavzevii*^78^, *S. paradoxus*^78^ and *S. uvarum*), and *S. chiloensis* sp. nov. x *S. uvarum* crosses (green dots). NRT = Non-Reciprocal Translocations and RT = Reciprocal Translocation. Spore viabilities from other crosses were obtained from Liti et al, 2006 ^78^. (D) Estimation of the divergence time since the last common ancestor between lineages. Time between (1) *S. eubayanus* and *S. uvarum*/*S. chiloensis* sp. nov. lineages; (2) *S. uvarum* and *S. chiloensis* sp. nov.; (3) the SA-C and AUS *S. chiloensis* sp. nov. lineages. Timeline is shown in Thousand’s years (Ky).

In addition, we assessed the presence of nuclear genome contributions from *S. uvarum* in *S. chiloensis* sp. nov. We detected introgression signals ranging from 2 – 79 kb on chromosomes V (*∼*70 kb), XI (*∼*49 kb) and XVI (*∼*19 kb) (**Figure S7, Table S11**). These findings conclusively establish the existence of a distinct lineage of *Saccharomyces* strains in coastal Patagonia and Tasmania, exhibiting significant divergence from the established *S. uvarum* lineages. The notable nucleotide divergence and OrthoANI values strongly suggest potential reproductive isolation, emphasizing the likelihood of these lineages representing a separate species.

Finally, we estimated the divergence time since the most recent common ancestor between lineages using the population genomic data as previously described in yeast^57,79^. The divergence time between *S. uvarum* and the SA-C and AUS lineages corresponds to approximately 473-59 KYA (**Figure 4D**), likely dating back to the Middle Pleistocene. On the other hand, the divergence between AUS and SA-C was dated to approximately 171-21 KYA, likely during the late Pleistocene (**Figure 4D**).

Our results indicate that *S. chiloensis* sp. nov. would represent a distinct *Saccharomyces* species. Additionally, the lack of many SVs between the AUS lineage and *S. uvarum* strongly supports that nucleotide divergence, rather than SVs, is responsible for the reduced spore viability between *S. chiloensis* sp. nov. and *S. uvarum*. In addition, the significant SV count between SA-C and AUS reinforces our observations of the lower offspring viability among these *S. chiloensis* sp. nov. lineages.

### Phenotypic diversity across *S. chiloensis* sp. nov. populations

***Description of Saccharomyces chiloensis sp. nov.*** Standard description of *Saccharomyces chiloensis* T. A. Peña, P. Villarreal, N. Agier, M. de Chiara, T. Barría, K. Urbina, C. A. Villarroel, A. R. O. Santos, C.A. Rosa, R. F. Nespolo, G. Liti, G. Fischer, and F. A. Cubillos sp. nov.

Etymology*: Saccharomyces chiloensis* (chi.lo.eńsis. N.L. f. adj. *chiloensis* of or pertaining to the Chiloé island (Chile), where this yeast was found).

On YM agar after 3 days at 25 °C, the cells are globose, ovoid or elongate, 2.8 -6.3 X 4.1 – 7.8 *μ*m, and occur singly or in small clusters (**Figure 5A**). In Dalmau plates after 2 weeks on cornmeal agar, pseudohyphae are either not formed or are rudimentary. Budding cells are transformed directly into persistent asci containing one to four globose ascospores formed after 11 days on YM agar at 25 °C (**Figure 5B**). Fermentation of glucose is positive. Assimilation of carbon compounds: positive for glucose, galactose, maltose (variable, weak/slow), sucrose, melibiose (weak/slow), raffinose, and D-manitol (variable, weak/slow). Negative for L-sorbose, cellobiose, trehalose, lactose, melezitose, inulin, soluble starch, D-xylose, L-arabinose, D-arabinose, D-ribose, L-rhamnose, ethanol, glycerol, erythritol, ribitol, galactitol, D-glucitol, salicin, DL-lactate, succinate, citrate, myo-inositol, methanol, hexadecane, xylitol, acetone, ethylacetate, 2-propanol, D-gluconate, and N-acetyl-D-glucosamine. Assimilation of nitrogen compounds: negative for lysine, nitrate, and nitrite. Growth in amino-acid-free medium is positive. Growth at 37 °C is negative. Growth in 1% acetic acid is negative. Growth on YM agar with 10 % sodium chloride is negative. Growth in 50 % glucose/yeast extract (0.5 %) is negative. Acid production is positive. Starch-like compounds are not produced. In 100 µg cycloheximide ml^-1^ growth is negative. Urease activity is negative. Diazonium Blue B reaction is negative (**Table S12)**. The CL1206 strain was designated as the type strain. The holotype of *S. chiloensis* sp. nov., strain CBS 18620^T^, is preserved in a metabolically inactive state at the CBS Yeast Collection of the Westerdijk Fungal Diversity Institute in Utrecht, the Netherlands. Additionally, two isotypes of *S. chiloensis* sp. nov., were deposited as strain RGM 3578, and PYCC 10014 in the Chilean Culture Collection of Microbial Genetic Resources (CChRGM) at the Agricultural Research Institute (INIA) in Chile, and the Portuguese Yeast Culture Collection in Portugal, respectively.

**Figure 5.**
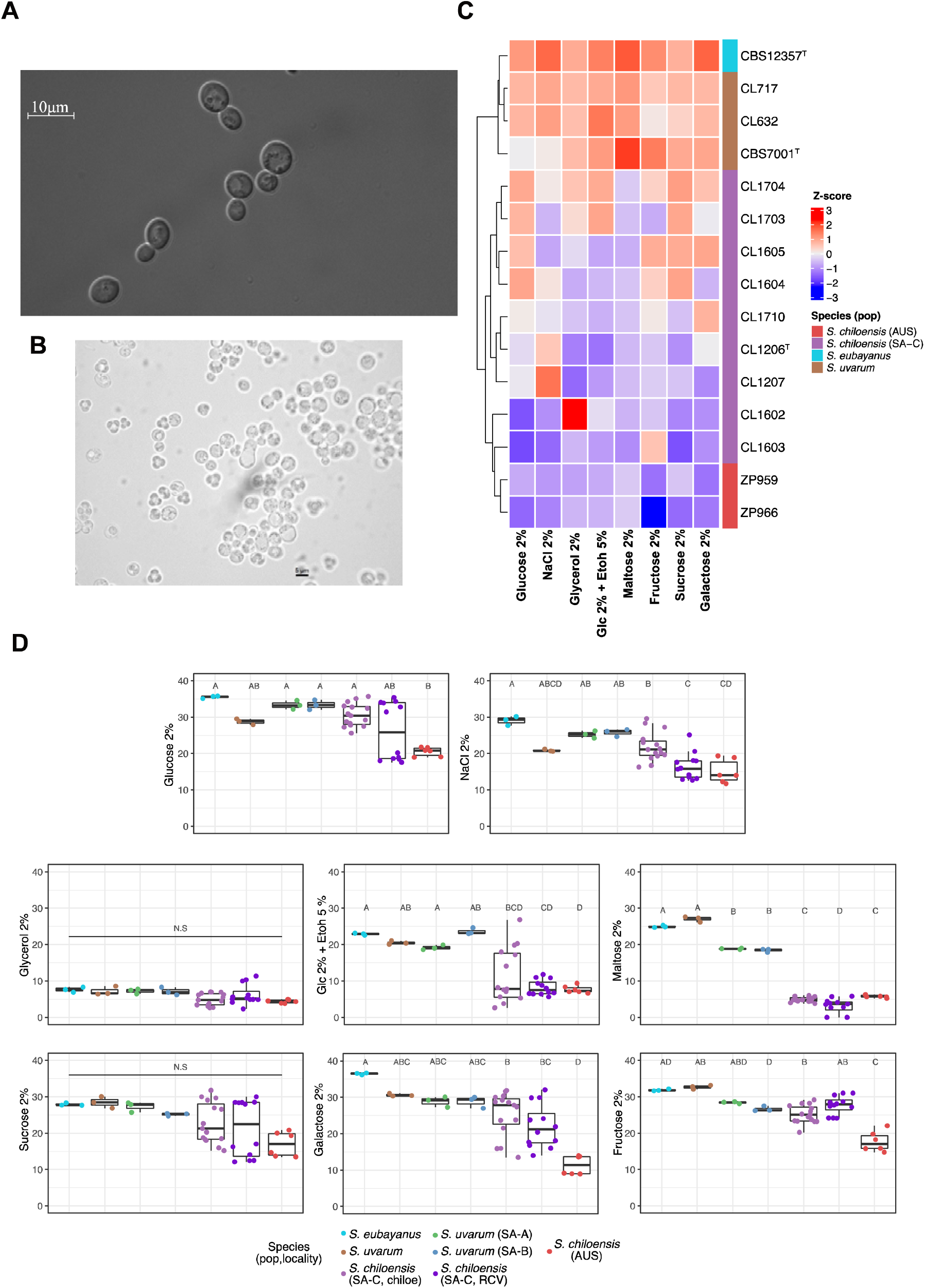
Phenotypic diversity in *S. chiloensis* sp. nov. (A) Differential interference contrast micrograph of budding cells of *S. chiloensis* sp. nov. grown in YPD broth after three days at 25°C. Bars: 10 *μ*m. This image was obtained using differential interference contrast (DIC) microscopy. (B) Budding cells and asci with ascospores on Yeast extract - Malt extract agar (YM) after three days at 25°C using 40X microscopy. (C) Heatmap depicting the phenotypic diversity in *S. chiloensis* sp. nov. obtained from 8 different environmental conditions. Strains are grouped by hierarchical clustering. The colours indicate the species and *S. chiloensis* sp. nov. (SA-C in purple, and AUS in red), *S. uvarum* (brown) and *S. eubayanus* (calypso). The heatmaps were obtained from the Area Under the Curve (AUC) data and normalized using Z-score per column (D) The Area Under the Curve (AUC) across all environmental conditions grouped by species, lineages and locality. Different letters reflect statistical differences between strains with a *P*-value < 0.05, one-way analysis of variance (ANOVA). Glc = glucose, EtOh = Ethanol.

Next, we explored phenotypic differences between *S. chiloensis* sp. nov. and *S. uvarum*. Through measurements of microbial growth in microcultures, we calculated the area under the curve (AUC) of growth curves under eight conditions that included different carbon sources, stresses, and environmental parameters (**Table S13**). Hierarchical clustering of the AUC data revealed two clades, clearly distinguishing *S. uvarum* and *S. eubayanus* strains from *S. chiloensis* sp. nov (**Figure 5C**). Notably, specific traits were identified as characteristic to *S. chiloensis* sp. nov., establishing differences from its sister species. We observed a significantly lower AUC in *S. chiloensis* sp. nov. strains grown under maltose and ethanol 5% relative to *S. uvarum (p*-value < 0,05, ANOVA, **Table S13**), suggesting a lower ethanol tolerance and capacity to assimilate carbon sources other than glucose and fructose. These findings highlight a distinct phenotypic profile in *S. chiloensis* sp. nov. relative to its sister species, *S. uvarum*.

Next, we sought to determine whether *S. chiloensis* sp. nov. exhibited intraspecies phenotypic variation. To investigate this, we compared the AUC among strains obtained from three different locations. Chiloé (SA-C-C), Reserva Costera Valdiviana (SA-C-RCV) and Australia (AUS). Interestingly, the AUS strains exhibited the lowest fitness across conditions compared to the other locations *(p*-value < 0,05, ANOVA, **(Table S13, Figure 5D**), particularly under galactose and higher maltose concentrations. These results indicate a separation of the isolate’s phenotypic profile according to their geographic origin, either from Patagonia or Australia, demonstrating substantial intraspecies phenotypic variation (**Figure 5C**).

## DISCUSSION

Species concepts and species delineation methods are essential when new lineages exhibit enough divergence to be declared a distinct species. However, the issue has been controversial due to the diversity of organisms’ lifestyles and reproductive modes^80^. For the case of microorganisms, the lack of fossil records for time’s calibration and their different reproductive modes, faces important challenges for new species identification, being a controversial topic for a long time^19,81^. Moreover, species delineation is essential to determining community variability and diversity^82^. Defining and classifying a microbial species mostly relies on utilizing a handful of markers, which may pose a significant challenge for correctly assigning phylogeny. For example, in many arbuscular mycorrhizal fungi, ITS markers were not resolutive enough relative to other nuclear markers^16,83,84^. When ITS and D1/D2 sequences provide insufficient phylogenetic precision, additional identification methods should be used. Generally, there is growing recognition that a single set of criteria does not adequately describe the diversity among fungal lineages^12,16,19,81^. An alternative is the utilization of integrative taxonomy, i.e., the combination of various species concepts: genome sequence-based classification, phenotype, and quantifying reproductive barriers^12,81^. Our study highlights the challenges associated with traditional species delineation methods in yeasts, mainly when relying on single genetic markers such as the ITS and LSU rRNA genes. Advances in genomics and the application of whole-genome data, specifically OANI have proven essential in overcoming these limitations^12,16,82^. The recognition of *S. chiloensis* sp. nov as a distinct evolutionary lineage, supported by OANI, emphasizes the significance of employing whole-genome-based approaches for accurate species delineation. Furthermore, mating tests, demonstrating reproductive isolation between both *S. chiloensis* sp. nov populations and *S. uvarum*, further support the identification of a taxonomically independent lineage. Soon, it would be interesting to assess the genomic compositions of surviving spores from interspecies crosses and determine whether viability was possible due to recombination events and postzygotic barriers between divergent genomes. A similar approach recently deciphered in *Cryptococcus* the genetic separation between *C. amylolentus*, *C. wingfieldii* and *C. floricola*, exhibiting ∼6% genetic divergence, significant chromosomal rearrangements and hybrid sterility^85^.

As revealed by our study, the evolutionary history of *Saccharomyces* species in Coastal Patagonia unveils a complex interplay of historical events and genetic differentiation (**Figure 6A**). The genetic relationship between *S. chiloensis* sp. nov and *S. uvarum*, hints at an allopatric speciation process, where a new species is generated due to a physical barrier. Intriguingly, the *S. chiloensis* sp. nov SA-C and AUS populations are found in Pacific islands located 42°S and associated with native *Nothofagus* forests, suggesting a common tree host with Gondwanan origin^86^. The restricted geographic distribution of the *S. chiloensis* sp. nov. populations on each side of the Pacific Ocean supports a secondary dispersal process driven by a vector, most likely bird. However, more evidence is needed to understand yeast dispersal across continents.

**Figure 6.**
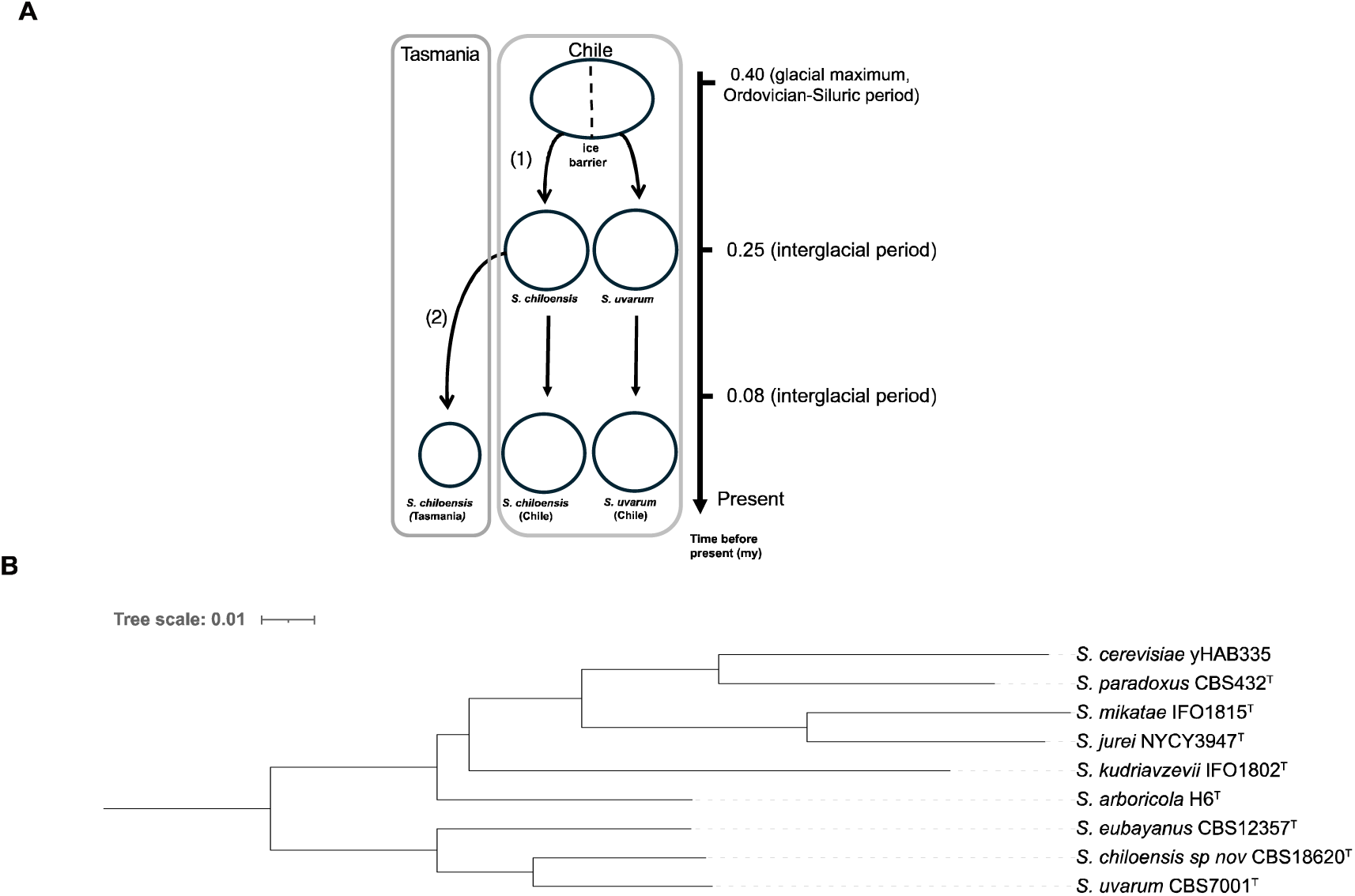
*Saccharomyces* species. (A) Hypothetical speciation-dispersal processes explaining the sympatric distribution of *S. chiloensis* sp. nov. and *S. uvarum*, together with the disjunct distribution of *S. chiloensis* sp. nov. in Chile and Tasmania. First, an ancestral *Sacharomyces* population experienced allopatric speciation (1) due to an ice barrier during the Middle Pleistocene glacial maximum (∼ 0.4 mya). Secondly *S. chiloensis* sp. nov. was dispersed by vectors to Tasmania (2), generating the actual distribution at present. (B) Maximum likelihood (ML) phylogenetic tree of the *Saccharomyces* genus using long-reads. The tree was built considering 6,615 orthologous genes using OrthoFinder. Branch lengths represent the average number of substitutions per site (tree scale).

In particular, *S. chiloensis* sp. nov is only found in glacial refugia during Pleistocene times, suggesting that this population did not expand after the glaciation period. Indeed, our analysis suggests that *S. chiloensis* sp. nov with *S. uvarum* split into two main clades during the middle Pleistocene (ca. 0.4-0.059 Ma), likely due to ice barriers during the recurrent glaciation periods in Patagonia. Later, the SA-C and AUS populations split during the Late Pleistocene. Similar findings were found between *Lachancea cidri* populations, estimating clades divergence between Australian and Patagonian lineages during the Late Pleistocene^10^. The potential impact of historical glaciation events on the genetic structure of *Saccharomyces* species in Patagonia, as inferred from shared ancestry and common glaciation periods, highlights the role of environmental factors and genetic drift in yeast evolutionary processes. The current sympatric existence of *S. chiloensis* sp. nov with *S. eubayanus* and *S. uvarum* in Patagonia suggests unique ecological niches and adaptive strategies related to these past events in the southern South Hemisphere. Future bioprospecting studies may expand our understanding of the genetic and ecological dynamics of *S. chiloensis* sp. nov, including its population structure and geographical distribution in the South Hemisphere.

The vast distribution of *Nothofagus* forests in Patagonia may promote a large genetic differentiation between *Saccharomyces* lineages, representing its main habitat in the South Hemisphere. In this sense, large genomic rearrangements, including chromosomal translocations and inversions, suggest that structural variations have shaped the genetic distinctiveness within the Patagonian *S. chiloensis* sp. nov population. The Australian population, on the other hand, does not show any major differences in structural variants from other *S. uvarum* populations. The presence of large structural variants is commonly observed within and between yeast species^78,79,87–91^. For example, *S. cerevisiae* strains exhibit a median of 240 SVs^91^, where massive genomic rearrangements have been reported in the Malaysian *S. cerevisiae* population, and a rapid accumulation of inversions and translocations in the South American *S. paradoxus* (formerly known as *Saccharomyces cariocanus*) that resulted in extensively altered genome configurations^89^. These large genomic rearrangements caused the partial reproductive isolation between these two lineages. However, in each case, the nucleotide divergence is below 1%; therefore, these lineages cannot be classified as distinct species. The reproductive isolation observed between *S. chiloensis* sp. nov and other *S. uvarum* lineages highlights the importance of considering both nucleotide divergence and structural variations in understanding the evolutionary trajectories of yeast species. Notably, the genetic divergence between *S. chiloensis* sp. nov and its closest relative*, S. uvarum*, exceeded that observed between other *Saccharomyces* populations, such as the average 4.6% sequence divergence between American and European *S. paradoxus* populations. This divergence and OANI values above the 95% threshold indicate that the two populations in *S. paradoxus* belong to the same species. On the contrary, *S. chiloensis* sp. nov exhibited nucleotide divergence greater than 6%, mating efficiency less than 6% and OANI values below the 95% suggested threshold relative to *S. uvarum*. In addition, we found that *S. chiloensis* sp. nov strains exhibited a distinct phenotypic divergence, marked mainly by differences in sugar consumption associated with carbon sources, such as maltose. These differences may correlate with the presence of structural variants in subtelomeric regions impacting the distribution of gene families associated with sugar consumption. These results demonstrate phenotypic differences between *S. chiloensis* sp. nov. and different lineages in *S. uvarum*. Interestingly, *S. chiloensis* sp. nov. strains exhibited a limited ability to grow under maltose conditions, suggesting that this species may have potential applications in producing low-alcohol beers. Additionally, investigating the metabolic pathways and regulatory mechanisms underlying this strain’s maltose utilization and aroma production could provide insights into utilizing this species for specific brewing applications. Altogether, our integrative taxonomic approach demonstrates that *S. chiloensis* sp. nov. represents a distinct evolutionary lineage.

Our findings emphasize the need for continued exploration and isolation of yeast strains to unravel the full extent of yeast diversity. As demonstrated in this study, advanced genomic tools and integrative taxonomy are crucial to uncovering hidden lineages and understanding the evolutionary forces shaping yeast populations. Future research should focus on expanding the repertoire of *S. chiloensis sp. nov.* strains from various geographic regions and habitats to precisely describe the genetic and phenotypic diversity within the species. Furthermore, similar genomic studies in diverse microbial systems can facilitate the identification of novel species and elucidate their roles and potential contributions to natural ecosystems and biotechnological applications. Understanding *Saccharomyces* species’ genetic and ecological dynamics, including the newly discovered *S. chiloensis* sp. nov, can have broader implications for biotechnological applications. Genetic diversity within *Saccharomyces* species, exemplified by distinct lineages in Coastal Patagonia, could offer valuable traits for industrial applications and explore its biotechnological potential, including its fermentation potential for low-alcohol beers. In conclusion, the discovery of *Saccharomyces chiloensis* sp. nov. in Coastal Patagonia expands our understanding of yeast diversity, challenges traditional species delineation methods, and highlights the importance of integrative taxonomy in unraveling the evolutionary dynamics of yeast species. The ecological and biotechnological implications of this discovery open new avenues for research in yeast biology and have broader implications for the fields of microbiology and biotechnology.

## Supporting information

Response to Reviewers

## ACKNOWLEDGMENTS

We acknowledge Fundación Ciencia & Vida for providing infrastructure, laboratory space, and experiment equipment. This research was partially supported by the supercomputing infrastructure of the National Laboratory for High Performance Computing Chile (NLHPC, ECM-02).

## AUTHOR CONTRIBUTIONS

**Conceptualization**: F.A.C and T.A.P; **Formal analysis**: F.A.C, T.A.P, P.V., N.A., G.F., M.dC., C.A.V., A.R.O.S., C.A.R., R.F.N., G.L. **Investigation**: T.A.P., P.V., N.A., T.B., K.U., A.R.O.S.: **Writing – original draft**: F.A.C and T.A.P. **Writing – review & editing**: F.A.C, T.A.P., P.V., G.F., G.L., R.F.N.

### Ethical statement

This article does not contain any studies with human nor animal subjects performed by any of the authors.

## SUPPLEMENTARY INFORMATION

### FIGURE LEGENDS

**Figure S1. Neighbor-joining phylograms of *RIP1* and *26S rRNA D1/D2 & ITS* markers.** (A) *RIP1* NJ phylogram. The tree was built based on the sequences of the RIP1 marker. The alignment was performed with MUSCLE. The distance metric is the number of substitutions. Bar, Substitutions per site. Bootstrap values are shown for each node. (B) Concatenated LSU-ITS-SSU NJ phylogram. The alignment was performed with MUSCLE. The distance metric is the number of substitutions. Bar, Substitutions per site. *S. jurei* sequences as outgroup. (C) Same as B but *S. eubayanus* as an outgroup.

**Figure S2. Admixture results.** Admixture plots (k=2 to k=10) for 100 individuals. Each color depicts a different lineage.

**Figure S3. Cross validation optimization.** For each k (1-10) the cross-validation error was estimated. A red square was shown to indicate the lowest value.

**Figure S4. FineSTRUCTURE Results.** The heatmap was obtained using fineSTRUCTURE chunk counts. Each row and column represent a strain, and the color scale indicates genetic sharing (yellow = low sharing, blue = high sharing). The tree shows the clusters inferred from the co-ancestry matrix. Populations and subpopulations can be inferred from the presence of darker colors in the diagonal. The strain matrix is orderly correlated. Below the figure colors represent lineages (brown: Hol, green: SA-A, blue: SA-B1, yellow: SA-B2, orange: SA-B3, Purple: SA-C and red: AUS).

**Figure S5. Pairwise nucleotide divergence between lineages.** The nucleotide divergence relative to the *S. uvarum* CBS7001^T^ strain is shown. Lineages and colors are depicted as follows: South America (SA-A (green), SA-B1 (blue), SA-B2 (yellow), SA-B3 (orange) and SA-C (purple)), Australia (AUS, red) and Holarctic (HOL, brown).

**Figure S6. Number of structural variants identified across lineages.** (A) Total number of Structural variants (SVs), (B) SVs larger than 1 kb, and (C) Divergence from alignment. Colors depict *S. chiloensis* sp. nov (AUS, red), *S. chiloensis* sp. nov (SA-C, purple), *S. eubayanus* (light blue) and *S. uvarum* (brown).

**Figure S7. Introgression analysis from *S. uvarum* on *S. chiloensis*.** Each plot represents the % divergence per site relative to *S. chiloensis* reference strain CBS18620^T^ on a 1 kb window (x-axis values are in bp). *S. uvarum* lineages are color-coded according to the key.

## TABLE LEGENDS

**Table S1a. Bioprospection and molecular identification of yeast isolates.**

**Table S1b. *S. uvarum* strains collection.**

**Table S1c. *S. uvarum* strains from previous studies.**

**Table S2a. Bioinformatics summary statistics from sequenced strains in this study.**

**Table S2b. Bioinformatics summary statistics from strains sequenced in other studies.**

**Table S3. *F_st_* results**

**Table S4. Estimated Nucleotide diversity on each population.**

**Table S5. Estimated genetic divergence between different populations and the *S. uvarum* reference genome (CBS7001^T^, Hol).**

**Table S6. Spore viability from crosses between lineages.**

**Table S7. Metadata and Metrics for genome assemblies using ONT long-read sequences.**

**Table S8. Set of structural variants detected between *S. chiloensis* sp. nov, *S. eubayanus* and *S. uvarum*.**

**Table S9. Size and number of genes involved in translocations events between *S. chiloensis* sp. nov and *S. uvarum*.**

**Table S10. OANI, nucleotide divergence and spore viability values in different *Saccharomyces* lineages and species**

**Table S11. Introgressions from *S. uvarum* on *S. chiloensis* sp. nov. CBS18620^T^.**

**Table S12. Phenotypic characterization of *S. chiloensis* sp. nov representative strain. Data from *S. eubayanus* (Libkind et al., 2011) and *S. uvarum* (Vaughan-Martini and Martini., 2011) is shown in gray.**

**Table S13. Phenotype data for *S. chiloensis* sp. nov strains obtained from growth curves analysis.**

