## Supplementary material for "An integrative taxonomy approach reveals Saccharomyces chiloensis sp. nov. as a newly discovered species from Coastal Patagonia": Response to Reviewers

**Manuscript number:** RC- 2024-02493

**Corresponding author(s):** Francisco, Cubillos

### 1. General Statements

### We want to thank the editors and reviewers for their time and effort in reviewing our manuscript, “*An integrative taxonomy approach reveals Saccharomyces chiloensis sp. nov. as a newly discovered species from Coastal Patagonia*.” We have now addressed all editorial and reviewers' comments and made the suggested changes to improve our manuscript.

### 2. *You will find a point-by-point response to the questions raised in the response to the reviewer’s file.*Point-by-point description of the revisions

Reviewer #1

This study by Peña et al. (Cubillos lab) provides multiple evidences of the existence of a highly divergent lineage of *Saccharomyces uvarum*, composed of strains isolated from Chilean Patagonia and Australia, which, according to the results obtained by the authors, can be considered as a new species (*S. chiloensis*).

*R: We would like to thank the reviewer for our manuscript's helpful and constructive comments. We have now addressed all comments and made the suggested changes.*

Major comments

I have no major comments on this work. They have use a broad array of genomic, phylogenetic and phenotypic data (including reproductive performance) to support their conclusion; in fact, I believe that all the claims of this work, including the main claim (delineation of a new species) are indeed supported by actual data.

R: *Thanks for the positive feedback on our strategy. Indeed, we tried to address the species delimitation challenge and cover the main conclusions by considering a wide array of strategies.*

I only want to suggest the inclusion of a code availability section, hopefully in a way that a well-structured and well-annotated script will make possible to easily connect raw data to final plots included in the main figures. I know that this suggestion will only be mandatory in certain journals, but it is a transparency effort that will make this work even more elegant.

*R: We have now included all the scripts and codes used in our study in the following Github: https://github.com/TomaasP/SchiloGenomics/tree/main*

Minor comments

I only have a minor experimental suggestion regarding the final phenotypic characterization of the species, which is assessing also the intra-specific diversity of those metabolic traits listed in Table S11, at least including 2-3 strains from each clade (AUS and SA-C). This could help to confirm those phenotypes currently defined as 'variable', and to help future researchers isolating new strains of this new species to better discuss unexpected results.

*R: We appreciate the reviewer's insightful suggestion regarding including a larger number of strains from the Australian population. While we agree that this would greatly benefit our study, we were limited to the two isolates kindly provided by Dr. Jose P. Sampaio at this study stage. We plan to expand our strain collection to include additional isolates from Chile, Australia, and New Zealand for future studies. We have now included an additional sentence in the discussion sections (lines 542-546) and it reads:*

*‘Future research should focus on expanding the repertoire of S. chiloensis sp. nov. strains from various geographic regions and habitats to precisely describe the genetic and phenotypic diversity within the species. Furthermore, similar genomic studies in diverse microbial systems can facilitate the identification of novel species and elucidate their roles and potential contributions to natural ecosystems and biotechnological applications.’*

I also suggest to include a new column in Table S1A, with the specific accession code (BioSample) for the genome of each isolate, just to make it easier to explore the dataset for future readers.

R: *This has been incorporated.*

Finally, I ask for further discussion on the potential industrial implications of this newly defined species, as it is mentioned by the authors several times throughout the manuscript. I do not see the specific industrial needs that can be covered by this new species, especially compared with other sister Saccharomyces species (i.e. S. uvarum, S. eubayanus, S. cerevisiae, etc).

R: *We agree with the reviewer's suggestion that we discuss the potential industrial applications of these strains. In response, we have included a new paragraph in the Discussion section addressing the potential use of these strains in producing low-alcohol beers. The added content can be found on lines 532—538, and it reads:*

*‘These results demonstrate phenotypic differences between S. chiloensis sp. nov. and different lineages in S. uvarum. Interestingly, S. chiloensis sp. nov. strains exhibited a limited ability to grow under maltose conditions, suggesting that this species may have potential applications in producing low-alcohol beers. Additionally, investigating the metabolic pathways and regulatory mechanisms underlying this strain's maltose utilization and aroma production could provide insights into utilizing this species for specific brewing applications. Altogether, our integrative taxonomic approach demonstrates that S. chiloensis sp. nov. represents a distinct evolutionary lineage.’*

**Referee Cross-commenting**

After reading the report from Reviewer #2, which aligns with my own report, I have no additional comments to add.

Reviewer #1 (Significance (Required)):

This work is of great importance for those interested in yeast biology, but also for the broader community of researchers studying global biodiversity patterns, especially in underexplored areas such as the global south. It provides a significant advance of knowledge in the field of yeasts biology and phylogeny, and its robust methodological design will guide future yeast bioprospecting effort that leads to the delineation of new species through an integrative taxonomy approach. The main limitation of this work is the limited number of isolates currently available for this species, and even for its sister species S. uvarum, which can limit the reliability of the population structure of the species, which could change in the future when several hundred isolates are available for study.

*R: We agree that our current study is based on a limited number of new isolates (44 S. chiloensis strains, 42 S. uvarum strains, and 43 previously published S. uvarum genomes). However, this study opens new possibilities for extending our findings and identifying a larger set of strains worldwide. We are currently working on a follow-up study that considers several experimental campaigns to increase the number of strains from the Southern Hemisphere, specifically from Chile, Australia, and New Zealand. This will allow us to compile a more comprehensive strain collection and further explore the biology of S. chiloensis. As a first approximation, the number of isolates considered in our current study is a good proxy for understanding the species' differentiation from sister species.*

*We would also like to highlight that our study considers the largest number of strains in a S. uvarum study to date, identifying novel populations and further clarifying the genetic structure of this species. We have included an additional sentence on future perspectives in the discussion section (lines 542-546), and it reads:*

*‘Future research should focus on expanding the repertoire of S. chiloensis sp. nov. strains from various geographic regions and habitats to precisely describe the genetic and phenotypic diversity within the species. Furthermore, similar genomic studies in diverse microbial systems can facilitate the identification of novel species and elucidate their roles and potential contributions to natural ecosystems and biotechnological applications. Understanding Saccharomyces species' genetic and ecological dynamics, including the newly discovered S. chiloensis sp. nov, can have broader implications for biotechnological applications.’*

Likewise, it would be advisable to study the prevalence of this new species in different habitats (other tree compartments, soils, commensal strains in animals, anthropic ecosystems and agro-industrial matrices, etc.) and studying its biogeographical distribution and the environmental determinants of its population abundance in nature (i.e.niche modelling), so that it would be easier to guide future isolation efforts. All this comments are just obvious limitations for a first study describing a new species, so I only want to suggest some future research lines.

R: *We agree with the reviewer that these questions are intriguing and exciting. As mentioned in our previous response, we are dedicating significant efforts to understanding the habitat of this novel species. We aim in the future to precisely identify the forests where it is present and explore their connection with historical events in Patagonia.*

Expertise of the reviewer: Yeast ecology, Microbial diversity patterns

-------------------------------------------------------------------------------------------------------------------------------

Reviewer #2 (Evidence, reproducibility and clarity (Required)):

This study reports the identification and characterization of a new yeast species Saccharomyces chiloensis. The assignment as a new species was based on whole genome analysis, genetic crosses, and phenotypic analysis. The work is largely well done. It wasn't clear to this reviewer if this is a homothallic species, and if so whether there is an HO gene and silent mating type cassettes.

R: *We would like to thank the reviewer for their positive feedback. Since we were able to sporulate the S. chiloensis strains and found viable spores from different tetrads, we identified these strains as homothallic. Genome annotations allowed us to identify the silent mating type cassettes, and this information is now available in the BioProject accession number PRJNA1084000.*

*We have included this information in the Results section (lines 304-305). This new sentence reads as follows:*

*‘Interestingly, diploid SA-C and AUS strains exhibited normal spore viabilities, demonstrating that they are homothallic and ruling out inherent reproductive issues (Table S6).’*

The nature of the hybrid spore mating products was also not entirely clear. There was a low frequency of spore germination, but some further analysis of the spores that do germinate is necessary. Are they fertile, and if so with whom.

R: *These strains are interspecies hybrids; therefore, we anticipated that spore germination rates would be low if they belonged to different species. We have now analyzed the nature of four F2 viable spores between S. chiloensis x S. uvarum. Interestingly, after 14 days, only 1 out of 4 spores could sporulate, suggesting infertility. The sole spore was infertile, and no viable meiotic products were found. We have incorporated these new results in the results section (lines 294- 297).*

‘*Crosses involving representative SA-C and AUS strains (CL1604 and ZP966 strains, respectively) exhibited markedly lower spore viability (6.02% and 5.09%, respectively, Table S6, Figure 2). The resulting spores were sterile, and several of them did not undergo tetrad formation.*’

Have their genomes been sequenced, and are they diploid, aneuploid, euploid?

R: *We decided not to assess the genomes of the few viable spores, as this was not one of the objectives of our study. Typically, the biological species concept requires assessing viability between strains, but not necessarily the genomic nature of the few viable spores. In the near future, it would be interesting to assess the genomic compositions of those few surviving spores and determine whether viability was possible due to recombination events between divergent genomes. We have included a perspective sentence in the discussion section (lines 471 - 475).*

*‘Furthermore, mating tests, demonstrating reproductive isolation between both S. chiloensis sp. nov populations and S. uvarum, further support the identification of a taxonomically independent lineage. Soon, it would be interesting to assess the genomic compositions of surviving spores from interspecies crosses and determine whether viability was possible due to recombination events and postzygotic barriers between divergent genomes.’*

Have the authors looked for regions of introgression across the genome of isolates of the novel species to see if there are regions that have introgressed from other closely related yeasts?

R: *We have now included a supplementary information (Figure S7, Table S11) depicting the introgressions from the sister species S. uvarum. We have also included a sentence in the results section (lines 361 – 363), and it reads:*

*‘In addition, we assessed the presence of nuclear genome contributions from S. uvarum in S. chiloensis sp. nov. We detected introgression signals ranging from 2 – 79 kb on chromosomes V (~70 kb), XI (~49 kb) and XVI (~19 kb) (Figure S7, Table S11).’*

A recent publication that the authors could cite and discuss involved similar detailed analyses establishing a new yeast species in Cryptococcus.

Genetic and Genomic Analyses Reveal Boundaries between Species Closely Related to Cryptococcus Pathogens.

Passer AR, Coelho MA, Billmyre RB, Nowrousian M, Mittelbach M, Yurkov AM, Averette AF, Cuomo CA, Sun S, Heitman J. mBio. 2019 Jun 11;10(3):e00764-19. doi: 10.1128/mBio.00764-19. PMID: 31186317

*R: This reference complements our findings very well. We have now included it in the Discussion section and discussed our results in the context of the findings in Cryptococcus (lines 473-477), and it reads:*

*‘Soon, it would be interesting to assess the genomic compositions of surviving spores from interspecies crosses and determine whether viability was possible due to recombination events and postzygotic barriers between divergent genomes. A similar approach recently deciphered in Cryptococcus the genetic separation between C. amylolentus, C. wingfieldii and C. floricola, exhibiting ∼6% genetic divergence, significant chromosomal rearrangements and hybrid sterility (Ryan Passer et al, 2019).’*

**Referee Cross-commenting**

I have read the cogent report from Reviewer #1 and think the suggestions and points raised are right on target, and overall well aligned with my own view. This is an interesting and exciting study that with revision will make an important contribution to the field.

R: *We appreciate the reviewer's insights. We hope that this revised version addresses all the suggested improvements to our manuscript.*

Reviewer #2 (Significance (Required)):

This is a strong study and one that will be of general interest given the discovery of a new yeast in the Saccharomycotina. The paper is interesting to read. It will appeal to fungal geneticists and genomicists, as well as those interested in discovering and characterizing new yeast species. The work is largely well done, but would benefit from revisions, as outlined above.

*R: We appreciate the reviewer's supportive comments.*
